# A Graph Neural Network Approach to Molecule Carcinogenicity Prediction

**DOI:** 10.1101/2021.11.10.468094

**Authors:** Philip Fradkin, Adamo Young, Lazar Atanackovic, Brendan Frey, Leo J. Lee, Bo Wang

## Abstract

Molecular carcinogenicity is a preventable cause of cancer, however, most experimental testing of molecular compounds is an expensive and time consuming process, making high throughput experimental approaches infeasible. In recent years, there has been substantial progress in machine learning techniques for molecular property prediction. In this work, we propose a model for carcinogenicity prediction, CONCERTO, which uses a graph transformer in conjunction with a molecular fingerprint representation, trained on multi-round mutagenicity and carcinogenicity objectives. To train and validate CONCERTO, we augment the training dataset with more informative labels and utilize a larger external validation dataset. Extensive experiments demonstrate that our model yields results superior to alternate approaches for molecular carcinogenicity prediction.

## 1 Introduction

Globally, cancer is the second leading cause of death, and accurate molecule carcinogenicity prediction holds promise in decreasing the likelihood of disease onset[1]. The prediction problem can be broken down by cause: cancer can be caused by random somatic mutations, exposure to harmful radiation, or molecule reactivity with DNA [2]. Two major methods for measuring chemical and DNA reactivity are carcinogenic and mutagenic experiments [3] [4]. Carcinogenic data is considered more accurate as it measures tumor growth in animals, however, experiments can be costly and time consuming. Mutagenicity experiments are conducted in bacterial cultures and tend to be significantly cheaper and faster, but have higher rates of false positives and negatives. *In silico* predictions using neural networks provide an appealing alternative, and can aid in experimental selection of compounds for costly carcinogenicity experiments.

Most carcinogenic compounds have been identified as a result of observational studies in sub-populations with increased cancer penetrance. This approach is effective in two scenarios: the first is if the compound is extremely carcinogenic, e.g. aristolochic acid, which was present in certain herbal supplements before being identified as one of the most potent compounds in the carcinogenic potency database (CPDB) [5]. The second is when a large enough sub population is repeatedly exposed to a moderately carcinogenic compound, such as chimney sweepers in England first identified by Percivall Pott in 1775 [6]. These approaches however, cannot identify chemicals of intermediate potency and prevalence, leading to continuous exposure of individuals to dangerous compounds. A high throughput computational method for predicting molecular carcinogenicity can provide a filter for identifying high likelihood carcinogenic compounds that would otherwise be missed with the traditional identification workflows.

Current state-of-the-art solutions for molecule carcinogenicity prediction train models using ASCII string representations of molecules through simplified molecular-input line-entry systems (SMILES) [7][8][9]. A major drawback of this approach is that the architecture of these tools does not make use of explicit molecular graphical structure represented by nodes (atoms) and edges (bonds). Graph neural networks (GNNs) are invariant to the ordering of atoms in a molecular graph and can leverage their respective node and edge features. Furthermore, existing works make use of molecular carcinogenicity datasets that are not sufficiently large, resulting in a high probability of over-fitting. Toxicology in the 21st Century Program (Tox21) and Toxicity Forecaster (ToxCast) are two other standardized chemical structure datasets used for toxicity prediction and include a small proportion of carcinogenic chemical compounds. However, Tox21 and ToxCast fall short in providing adequate measures of carcinogenicity as cancer characteristics like immunosuprescense and genomic instability are not included in their chemical bioassays [10].

In this work, we propose a novel system CONCERTO: Carcinogenicity prediction with graph neural networks, with the following three main contributions:

- To the best of our knolwedge, we are the first to use GNN approaches to identify carcinogenic molecules.
- We augment an existing carcinogenicity dataset with more informative labels and make use of a larger dataset for external validation.
- We explore the utility of transfer learning from large models, and multi-round pre-training on lower quality mutagenicity experiments to improve predictive capacity.

## 2 Related Work

In general, molecule carcinogenicity prediction remains a challenging problem. Zhang *et al*. introduced CarcinoPred-EL, an ensemble-based approach for predicting the carcinogenicity of chemicals using molecular fingerprints, achieving a relatively high test accuracy on a limited dataset [8]. Recently, Wang *et al*. expanded on existing methods and presented a neural network model (CapsCarcino), however, the model is not publicly available [7]. In addition, CapsCarcino did not leverage the naturally occurring graphical representation of molecular data, and the validation dataset does not provide opportunity to demonstrate statistically significant improvements due to the constrained size. Consequently, there is reason to believe that GNN models could lead to improved results for carcinogenicity prediction.

GNN models are a family of neural networks that are suited to graph-type input data that can enforce notions of node permutation invariance and equivariance [11, 12]. Recent advancements in the domain of GNN research have demonstrated success for the task of molecular property prediction [13] [14] [15] and drug discovery [16]. Duvenaud et al showed utility of graph convolutional network (GCN) as an alternative way of representing a molecular profile, analogous to molecular fingerprints [13]. In a subsequent work, Gilmer et al proposed message passing neural networks (MPNNs) to predict quantum properties of organic molecular compounds [14], later improved upon and extended by Yang et al [15]. In addition, Stoke et al have demonstrated the promising outcomes of the MPNN model in the domain of drug discovery, uncovering the previously unknown antimicrobial molecular compound *Halicin* [16]. Recently, Ying *et al*. have introduced an effective position encoding technique for transformer architectures, finding success in molecular problem domains [17]. Motivated by this opportunity and the recent success of GNNs in molecular property prediction, we propose the use of a GNN-based models for molecule carcinogenicity prediction [7].

## 3 Methods

### 3.1 Problem Formulation

At it’s core carcinogenicity prediction is a graph level regression problem: given a molecular representation the algorithm predicts an associated carcinogenicity value indicative of DNA reactivity. We consider two types of molecular representations the first of which are graph based models, such as GNNs, use molecular graph representation in which every node represents an atom in the molecule, with edges indicating bonds between atoms. The node and edge features depend on the choice of model, but include important chemical properties like atomic number, atomic mass, and bond order. The second, fingerprint-based models use hand engineered features that aim to summarize important molecular properties like molecule aromaticity, presence of functional groups, and atom co-occurrence [18][19]. Fingerprints are an effective way to incorporate domain expert knowledge through identification of important molecular substructures.

A fundamental technical challenge at the core of carcinogenicity prediction problem is the difficulty of data acquisition. Every experiment is conducted in animals, usually rodents, resulting in dozens of animals required for a single data point. Not only does this result in limited training data, but presents difficulties for robust model evaluation. In addition carcinogen dosing is modulated using the maximum tolerated dose (MTD) where animals are given the highest possible compound dose without compromising animal survival. This leads to unrealistic quantities of chemical exposure, inflating the amount of tumor growth [5]. This results in datasets containing molecules labeled as carcinogenic, however having multiple orders of magnitude difference in their dosage potency. An effective relevant model would be able to distinguish molecules between dosage extremes. In certain experiments it is possible to express results through a dose-rate formulation represented by TD50 mg/kg body weight/day, which captures the differences of compound dosage as a proportion of body weight.

### 3.2 CONCERTO

To this end we propose CONCERTO: carcinogenicity prediction with graph neural networks (Figure 1). The method consists of 3 main parts: a large self-supervised graph neural network transformer, a multilayer perceptron (MLP) optimized over a molecular fingerprint representation concatenated with GNN transformer representation, and iterative fine tuning using multi-round pre-training on mutagenicity and carcinogenicity objectives.

**Figure 1:**
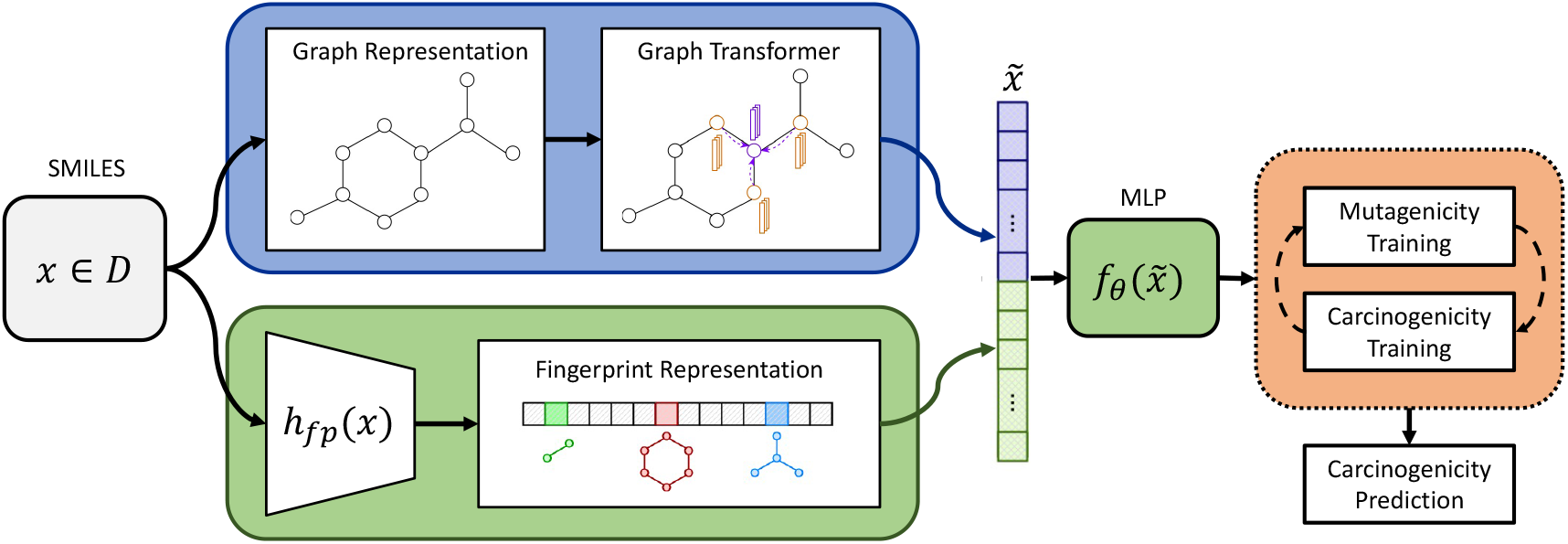
CONCERTO consists of two architectural parts. The first (in blue) is the GNN transformer, which takes in the graph representation of the molecule. The second (in green) is the predictor which consists of fingerprint representation of the molecule that is fed into the multilayer perceptron along with the GNN representation. The two models are jointly optimized with multi-round pre-training to generate carcinogenicity prediction.

#### 3.2.1 Graph Neural Net Transformer

In this section we provide background on GNNs, the attention operation, and finally the specifics of transfer learning from the GNN transformer [20].

First, given of a set of nodes (or vertices) 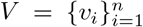 and a corresponding set of edges *E* = {(*v*_*i*_, *v*_*j*_)| *v*_*i*_, *v*_*j*_ ∈ *V, i, j* = 1, …, *n*}, a graph is defined as a tuple *G* = (*V, E*) of the respective node and edge sets. GNNs take the graph structure *G* as input in the form of an adjacency matrix and use node-wise and/or edge-wise layer embeddings to learn a non-linear predictive mapping. One such GNN is the MPNN which aggregates information in the form of “messages” across neighbourhoods of respective nodes [14]. For a one-hop neighbourhood, the hidden states of the *i*^*th*^ node of a MPNN are calculated as

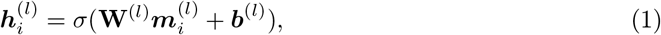

where ***W*** ^(*l*)^ are the neural network weights, *σ*(·) is some non-linear activation function, 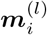 are the aggregated messages on the *i*^*th*^ node, and *l* is the message passing iteration number.

Second, the attention operation is fundamental to the transformer architecture, where an attention weight is computed for every pair of adjacent nodes using a linear embedding of node features. For the case of multi-head attention, given a set of queries ***q***, keys ***k***, and values ***v***, the attention operation is defined as

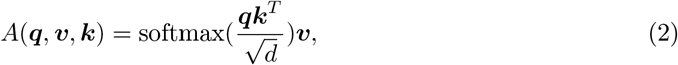

where *d* is the dimensionality of ***q*** and ***k***. The GNN transformer leverages a MPNN to embed the input data, presented as graphs, as query, key, and value vectors which are then fed into a multi-head attention network.

For each molecule we generate a representation using the 100 million parameter GROVER: graph representation from self - supervised message passing transformer [20]. It is a large graph neural network that is pre-trained on 10 million unlabeled molecules utilizing contextual and graph level motif prediction tasks. Contextual property prediction task consists of selecting a molecular subgraph and using it’s representation to predict local neighborhood properties. Motif prediction task involves an alternative way to encode fingerprint representations in a structure: the task consists of predicting presence of a functional group given the entire molecular representation. Self-supervised training is conducted over representations of nodes and edges, making use of the attention operation to aggregate information across local neighborhoods. During training we freeze GNN transformer weights and use the computed representation as input to the MLP [20].

#### 3.2.2 Multilayer Perceptron Fingerprint Predictor

To add explicit structure information we supplement the implicit representation learned from the graph transformer with fingerprint data. We encode each molecule using Morgan, RDKit and MACCS fingerprints to capture properties relating to molecule aromaticity and functional groups. [21][19][18][22]. We concatenate the representation from GNN transformer to the fingerprints and train a multilayer perceptron to predict molecular carcinogenicity. We utilize ReLU activations and batch normalization to stabilize training.

#### 3.2.3 Multi-Round Mutagenicity Pre-training

To improve carcinogenicity model predictions, we pre-train on related, lower accuracy, but abundant mutagenicity experiments. Instead of measuring tumor growth in animal systems to evaluate carcinogenicity, mutagenicity experiments measure compound DNA reactivity in cellular systems. DNA damage is usually evaluated with an indirect measure such as cell growth resulting in noisy measurements with lower rates of reproducibility [23]. Although lower quality, mutagenicity experiments are an order of magnitude more abundant and measure a related property to carcinogenicity. We find that multi-round pre-training increases model performance for carcinogenicity prediction.

For mutagenicity pre-training and evaluation we use a dataset generated by Hansen *et al*. (Table 1) [24]. There is a significant overlap between mutagenicity and carcinogenicity datasets, but limited concordance: only 70%. The high mismatch percentage is in part due to low reproducibility of mutagenicity experiments as a result of simplified experimental systems [23][25]. To identify overlapping molecules we standardize molecular SMILES representation, since there is a one to many mapping between a unique molecule and SMILES strings. Missing this crucial step can lead to an overlap between training and validation sets resulting in inflated performance [26].

**Table 1:** Summary statistics of chemical compound carcinogenicity datasets. Under *Experiment Type*: C stands for carcinogenic experiments, M stands for mutagenic experiments. A significant fraction of compounds are present in multiple databases.

| Dataset | Experiment Type | # of Experiments | (+) Labels | (-) Labels |
| --- | --- | --- | --- | --- |
| CPDB | C | 6540 | 509 | 494 |
| CCRIS | C+M | 88056 | 2674 | 2099 |
| Hansen | M | N/A | 3403 | 2909 |

#### 3.2.4 Model Selection

We perform a hyper-parameter sweep over model architecture features, training parameters, and pre-training parameters. For each set of hyperparameters we perform 3-fold cross validation and rank the models based on the average validation performance of the folds. We then choose the top 3 performing models, average their prediction and evaluate on the test sets.

### 3.3 Counterfactual Approach to Model Interpretability

Exmol [27] is a model-agnostic method for interpreting chemical property predictions. To investigate a particular prediction, Exmol creates a local chemical subspace around the target molecule and searches for nearby counterfactual examples. The subspace is generated by randomly mutating the SELFIES [28] string representation of the target [29]. After applying the model to each molecule in the subspace, Exmol can find examples that are chemically close to the target with drastically different predictions. These counterfactual molecules can help explain the model’s behaviour by highlighting differences in the input (i.e. functional groups, rings) that have a large effect on the output.

## 4 Experimental Design

In this work, we consider a selection of chemical compound databases comprised of long-term carcinogenesis bioassays in animals, as well as short term mutagenicity experiments in bacterial cultures. To tackle the MTD problem we augment our training set and use transformed TD50 values instead. In addition we assemble a new external test set for evaluating carcinogenicity 5 times larger than the previous [30]. To confirm the validity of the new carcinogenicity test set we evaluate the distances between molecular distributions using Tanimoto scores.

### 4.1 Continuous Carcinogenicity Measure

For model training we utilize CPDB, which is a collection of 6540 experimental tests containing results from long-term carcinogenisis bioassays, primarily in rodents, for over 1000 chemical compounds (Table 1) [5]. Carcinogenicity of the tested chemical compounds was determined using TD50 values, an estimated numerical measure of carcinogenic potency, which represent the dose-rate of tumor development. Instead of using binarized tumor growth labels we use log reciprocal TD50 values for model training, yielding a richer information label. TD50 is estimated using the proportional hazards model[31][32]:

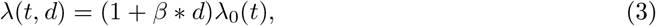

where *λ*_0_(*t*) is the tumor incidence at no dosing and *β* = 0 when there is no relationship between molecular dosage and tumor growth [33]. While TD50 is calculated in the following way *log*(2)/*β*, we instead transform the score to be *log*(*β*) for numerical stability during training. In addition, to summarize data across multiple experiments we take the harmonic mean over TD50 values, which biases the results towards low values, i.e. experiments that demonstrated molecular carcinogenicity. Our reasoning follows that of the CPDB authors’: given that a single experiment demonstrated carcinogenicity, the compound is likely to have some carcinogenic properties that are present in a unique set of conditions.

### 4.2 External Test Set

For external test evaluation we use CCRIS, which is a database containing experimental test results of over 4500 chemical compounds gathered from various studies cited in literature [34]. These experiments were conducted on chronic cancer animal models, the majority of which were rodents, measuring carcinogenicity, mutagenicity, tumor promotion, and tumor inhibition. A panel of experts used the aggregated experimental results to assess the molecular carcinogenic and mutagenic labels. To our best knowledge no works have used CCRIS for evaluating or training classifiers predictive of molecular carcinogenicity.

### 4.3 Estimating Differences of Molecular Distributions

In this work we introduce a new dataset for carcinogenicity analysis and use a perturbation approach for evaluating functional group importance, underscoring the utility of measuring molecular distances. We use Tanimoto molecular similarity as a measure for inter-molecular proximity, molecular diversity to measure dataset variance, and maximum mean discrepancy (MMD) to measure dataset distances [35].

To calculate Tanimoto similarity, we draw two molecules from their respective datasets 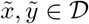 and calculate the Morgan binary fingerprints [18] as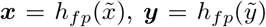, where *h*_*fp*_ is a mapping from SMILES strings to vectorized binary representations. Then, we define the Tanimoto similarity coefficient as

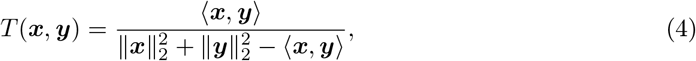

where ⟨·, ·⟩ is the vector dot product and ⟩ · ⟩_2_ the Euclidean norm [36].

Diversity is a metric used to quantify molecular variability [37] [38]. Given data 𝒳 of molecular fingerprints and the Tanimoto similarity coefficient *T* (***x, y***), we define the diversity score between any two samples ***x***_*i*_, ***x***_*j*_ ∈ 𝒳 as

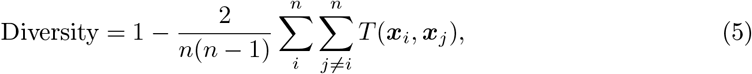

where *n* = |𝒳|.

We make use of the MMD score to define a distance metric between molecular datasets. Given two sets of molecular fingerprints that are sampled from two distributions 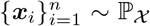 and 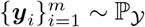 and for some similarity kernel function *k*(***x***_*i*_, ***y***_*i*_), the empirical estimate for MMD is defined as

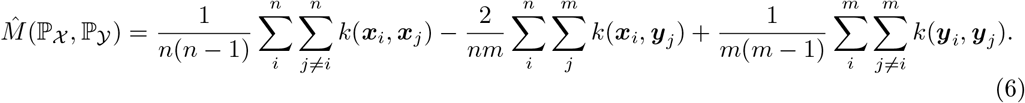

We use the Tanimoto similarity coefficient as the kernel function, i.e. *k*(***x***_*i*_, ***y***_*i*_) = *T*(***x***_*i*_, ***y***_*j*_).

## 5 Results

### 5.1 Model results

We perform hyperparameter searches over CONCERTO architectures, fingerprint baseline, and standalone GNN transformer, and find that CONCERTO outperforms the baseline models on the both test sets (Table 2). Standalone GNN transformer results in high variability predictions underscoring the importance of the MLP-fingerprint model component. Most importantly we observe that CONCERTO outperforms other models in the low false positive region, the prediction regime in which the carcinogenicity compounds can be identified with the lowest false discovery rate (Figure 2). Previous state-of-the-art, CarcinoPred-EL, was trained on data from CPDB, therefore we are unable to generate predictions without overfitting. Instead, we compare on an external CCRIS test set where our model outperforms all variants of CarcinoPred-EL (Table 2). We find that MLP-fingerprint predictor in conjunction with GNN transformer deliver the best results on the CPDB test set. We hypothesize that in data constrained settings, fingerprints are an effective way for representing domain expert knowledge.

**Table 2:** Performance of CONCERTO, baselines, and CarcinoPred-EL on CPDB and CCRIS. ROC, and PR values accompany plots a, b from Figure 2 and are calculated only over values for which CarcinoPred-EL is defined for. CarcinoPred-EL was trained on CPDB so we are unable to generate predictions without confounding overfitting. Uncertainty is calculated using standard deviation over data re-sampled with replacement (bootstrapping).

| Model | CPDB<br>n=518 | CPDB<br>n=518 | CCRIS<br>n=202 | CCRIS<br>n=202 |
| --- | --- | --- | --- | --- |
|  | Pearson | MSE | ROC AUC | PR AUC |
| CONCERTO | **$0.50 \pm 0.04$** | **$0.71 \pm 0.06$** | **$0.73 \pm 0.03$** | **$0.72 \pm 0.01$** |
| Fingerprint Baseline | $0.36 \pm 0.07$ | $0.81 \pm 0.08$ | $0.68 \pm 0.03$ | $0.64 \pm 0.01$ |
| GNN Transformer Baseline | $0.15 \pm 0.16$ | $0.83 \pm 0.07$ | $0.68 \pm 0.10$ | $0.69 \pm 0.01$ |
| CarcinoPred-EL Average RF | - | - | $0.67 \pm 0.02$ | $0.65 \pm 0.02$ |
| CarcinoPred-EL Pubchem RF | - | - | $0.64 \pm 0.03$ | $0.61 \pm 0.01$ |

**Figure 2:**
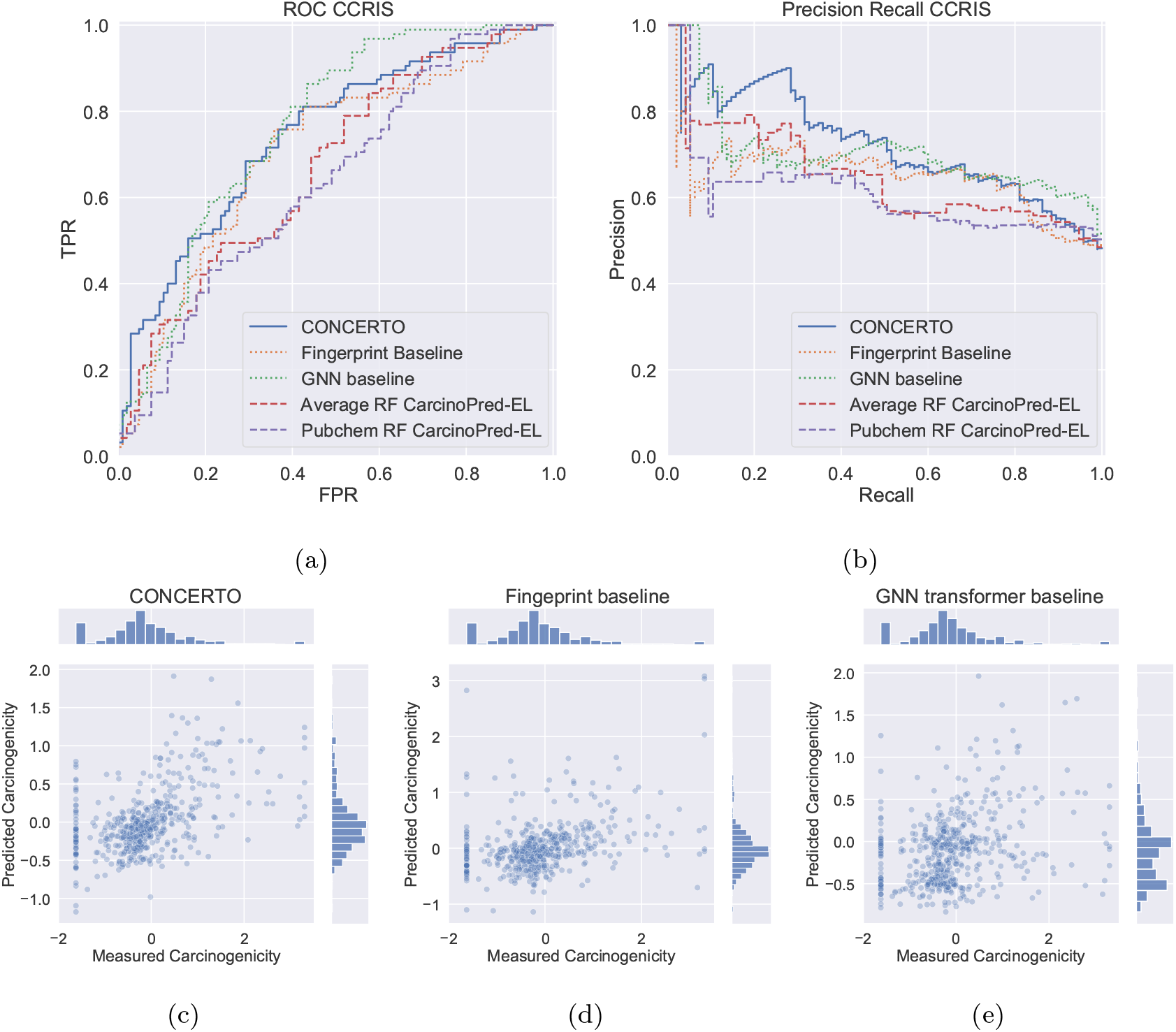
(a,b) ROC and Precision-Recall plots demonstrating performance gains of CONCERTO (solid) over previous state of the art (dashed) on an external test dataset, CCRIS. (c-e) Correlation between log reciprocal TD50 values and model predictions on the CPDB test set. A clustered set of points at -1.62 carcinogenicity values indicates experiments in which no tumor growth was observed in animals.

### 5.2 Quantifying dataset differences

To better understand differences between our datasets, we calculate maximum mean discrepancy over Tanimoto scores, as described in equations 4, 6. We perform MMD calculations for carcinogenic datasets while further partitioning data into positive and negative classes (Figure 3). Our first observation is that as expected, the distance between matching classes across datasets is smaller than the within dataset distances between positive and negative classes. This indicates that the inter-dataset differences are smaller than inter-class differences, confirming our choice of using CCRIS as an external test dataset. Our second observation is that inter-class distances vary between different datasets. We observe that CCRIS MMD distance between positive and negative classes (0.024) is significantly greater than CPDB inter-class distance (0.009) which leads us to hypothesize about the differing nature of dataset construction. One reason for this observation could be due to the fact that CCRIS labels were assigned by a panel of experts with molecules selected at either end of the carcinogenicity spectrum. Meanwhile CPDB inclusion criteria consisted of a robust set of experimental criteria followed by calculating a TD50 score. These observations support our choice for utilizing CCRIS as an external test set for CPDB trained models.

**Figure 3:**
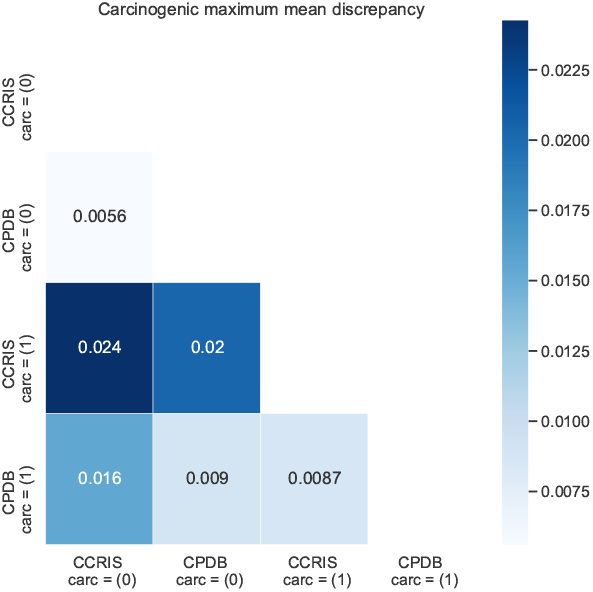
Analysis of dataset distances are generated by calculating MMD over Tanimoto scores. For carcinogenicity there are two datasets, that we further subdivide into positive and negative labels, creating four partitions. For each pair of partitions we calculate corresponding distances and visualize using a heatmap.

### 5.3 Ablation experiments

We perform experiments to identify individual contributions to changes in performance of mutagenicity pre-training and addition of GNN transformer representation. We conduct ten runs with matched seeds on a well performing set of hyperparameters to evaluate whether addition of pre-training improves performance on CPDB and CCRIS. We find that providing GNN transformer representation to the models improves performance on both the test sets (Table 3). The mutagenicity pre-training objective further improves performance almost matching the performance of the full CONCERTO model: a combination of GNN transformer and multi-round mutagenicity pre-training.

**Table 3:** Ablation experiments for CONCERTO models measuring the impact of graph neural network transformer, and multi-round mutagenicity pre-training. All architectures contain the MLP-fingerprint predictor. Results are averaged over 10 random seed runs. Standard deviation is computed over the random seed results.

| Experiment | CPDB correlation | CCRIS ROC |
| --- | --- | --- |
| 3 Mutagenicity pre-training + GNN | $0.36 \pm 0.03$ | $0.73 \pm 0.02$ |
| 1 Mutagenicity pre-training + GNN | $0.35 \pm 0.03$ | $0.71 \pm 0.02$ |
| No mutagenicity pre-training + GNN | $0.30 \pm 0.05$ | $0.70 \pm 0.02$ |
| No mutagenicity pre-training + No GNN | $0.21 \pm 0.09$ | $0.60 \pm 0.04$ |

### 5.4 Counterfactuals for Model Interpretability

We utilise Exmol to identify molecular substructures that are important drivers in carcinogenicity prediction [27]. In Figure 4 we demonstrate a method for model interpretability in which molecular substructures are added or removed from the original molecule. These changed molecules are counterfactual examples that are close to the original as measured by Tanimoto distance, however have large changes in carcinogenicity predictions. In the two demonstrated examples, replacing iodine and nitrogen atoms leads to decreased predicted carcinogenicity allowing us to speculate about their reactive nature. Similarly, in the increased carcinogenicity counterfactual molecules, there is an addition of a nitrogen and sulfur atom.

**Figure 4:**
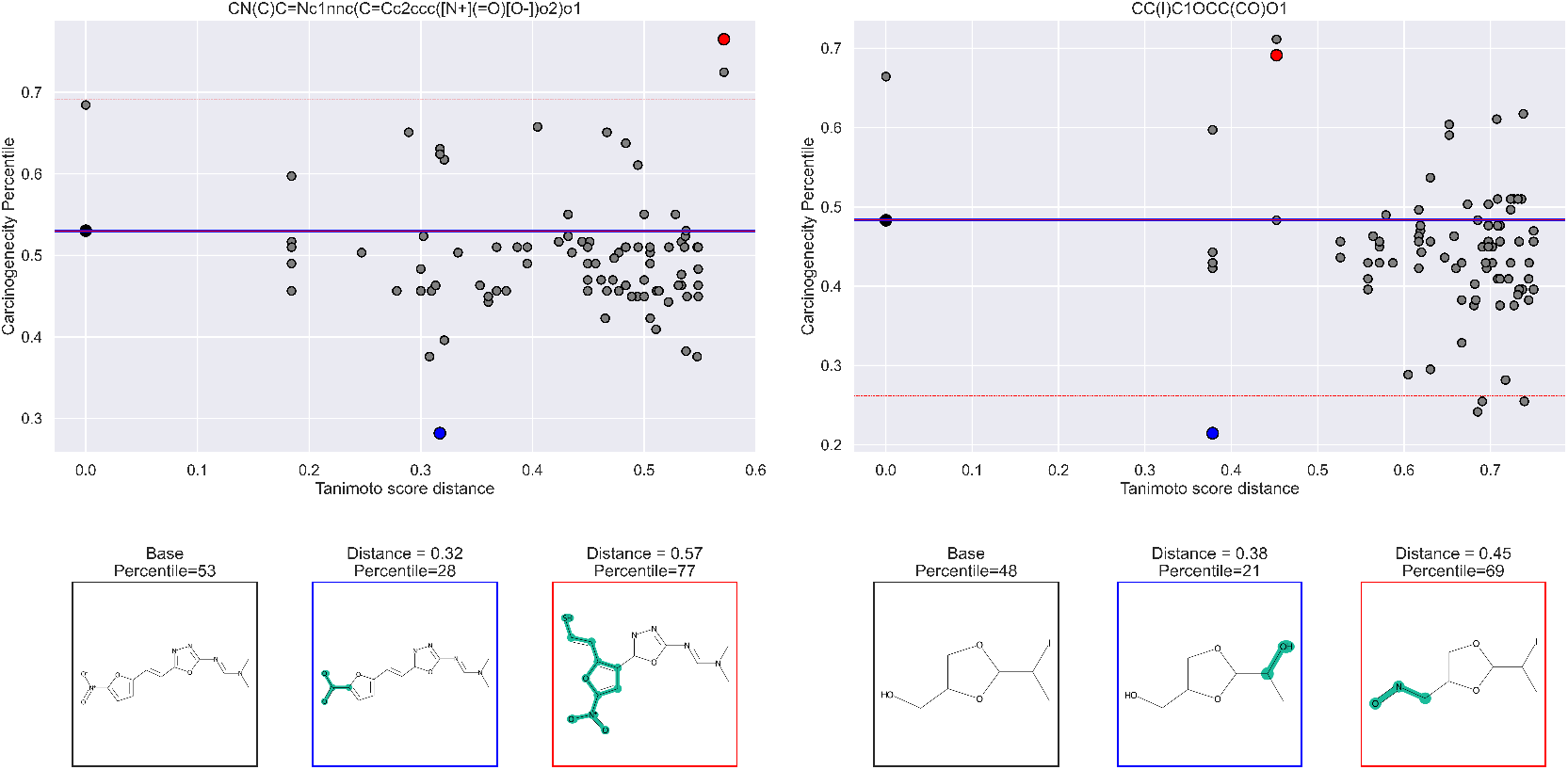
Two examples of a counterfactual analysis. On the x axis we’re plotting Tanimoto score distances between mutated molecules and the original molecule. On the y axis we’re plotting the predicted carcinogenicity relative to test set carcinogenicity distribution. For each molecule we visualize a positive and negative counterfactual examples. Average within dataset diversity as measured by Tanimoto distances is 0.88.

## 6 Discussion

Predicting molecular carcinogenicity is an important public health problem to address, however due the prohibitive cost and difficulty of measuring carcinogenicity, many compounds lack experimental data. In this work, we investigate three orthogonal approaches to overcome dataset size constraints: architecture choice, dataset modification, and pre-training techniques. First, we leverage the inductive bias present in GNN architectures relating to the graphical molecular structure. We find that the GNN transformer in conjunction with a MLP-fingerprint predictor outperforms the fingerprint baseline as well as the previous state-of-the-art. We suspect that due to the limited size of available data, a combination of a fine tuning a large GNN, and hand-engineered features extracted from molecular structures, is effective at capturing important drivers of molecular carcinogenicity. Next, we augmented the dataset with more informative labels by aggregating individual experimental results and creating continuous labels. This creates a richer representation for the network and circumvents the MTD design problem, where a molecule could be carcinogenic at a maximum dose for the animal but it would be impossible to be exposed to that dosage in the natural world. In addition, we collected an external dataset five times larger than previous, allowing us to make meaningful model performance comparisons while decreasing concern of overfitting to the test set. Finally, we explore the utility of model pre-training in two forms: first utilizing transfer learning from a large GNN transformer model, and second multi-round pre-training on related but lower quality experiments. We make use of a self-supervised GNN transformer embedding and fine tune it on our task of interest. Although during pre-training the transformer GNN was not shown any carcinogenicity data, through the self-supervised property and motif prediction tasks it was able to learn molecular properties that are predictive of carcinogenicity. In addition, we find that multiple rounds of mutagenicity pre-training significantly improves model performance. The differentiable nature of our model allows us to make use of effective pre-training strategies. We find that out of the three approaches, pre-training strategies resulted in the largest increase in performance.

We hope that this model will aid in identifying molecules for downstream carcinogenicity prediction as well as analysis of molecular perturbations resulting in decreased carcinogenicity. Given that up to 13% of recent drug retractions have been due to molecular DNA reactivity, a method for identifying functionally similar molecules but with decreased carcinogenicity could be useful [39]. To that end, we demonstrate an approach for visualizing counterfactual examples (Figure 4). We aim for that this technique being useful to domain experts for interpreting model predictions and iterating on the molecular design process.

## 7 Conclusions

In this work we present a model trained to predict molecule carcinogenicity, a dataset for validating model performance, and a counterfactual method for model interpretability. Our model, CONCERTO, makes use of transfer learning from a GNN transformer, multi-round mutagenicity pre-training, and continuous labels for addressing the maximum tolerable dose problem. We find that the combination of these training techniques results in a model that outperforms the previous state-of-the-art. Our code is publicly accessible and can be found here.

